# Analyzing hCov Genome Sequences: Predicting Virulence and Mutation

**DOI:** 10.1101/2020.06.03.131987

**Authors:** Shashata Sawmya, Arpita Saha, Sadia Tasnim, Md. Toufikuzzaman, Naser Anjum, Ali Haisam Muhammad Rafid, M. Saifur Rahman, M. Sohel Rahman

**Author notes:** Equal contributions (Shashata Sawmya, Arpita Saha, Sadia Tasnim).

## Abstract

**Background:** Covid-19 pandemic, caused by the SARS-CoV-2 genome sequence of coronavirus, has affected millions of people all over the world and taken thousands of lives. It is of utmost importance that the character of this deadly virus be studied and its nature be analyzed.

**Methods:** We present here an analysis pipeline comprising a classification exercise to identify the virulence of the genome sequences and extraction of important features from its genetic material that are used subsequently to predict mutation at those interesting sites using deep learning techniques.

**Results:** We have classified the SARS-CoV-2 genome sequences with high accuracy and predicted the mutations in the sites of Interest.

**Conclusions:** In a nutshell, we have prepared an analysis pipeline for hCov genome sequences leveraging the power of machine intelligence and uncovered what remained apparently shrouded by raw data.

## 1 Introduction

Covid-19 was declared a global health pandemic on March 11, 2020 [1]. It is the biggest public health concern of this century [2]. It has already surpassed the previous two outbreaks due to the coronavirus, namely, Severe Acute Respiratory Syndrome Coronavirus (SARS-Cov) and Middle East Respiratory Syndrome Coronavirus (MERS-Cov). The virus acting behind this epidemic is known as Severe Acute Respiratory Syndrome Coronavirus 2 or SARS-CoV-2 virus, in short. It is a single stranded RNA virus which is mainly 26,000 to 32,000 bases long on average [3]. The novel coronavirus is spherical in shape and has spike protein protruding from its surface. These spikes assimilate into human cells, then undergo a structural change that allows the viral membrane to fuse with the cell membrane. The host cell is then attacked by the viral gene through intrusion and it copies itself within the host cell, producing multiple new viruses [4].

The GISAID initiative database [5] has been collecting high quality complete genome sequences of the SARS-CoV-2 virus from clinicians and researchers from around the world since the beginning of the COVID-19 outbreak. To understand the virulence of the genome sequences and the nature of viral mutation, here, we present an analysis pipeline of the genome sequence leveraging the power of machine intelligence.

This paper makes the following key contributions:

1. Several Machine Learning and Deep learning models are used to identify the virulence of the genome sequences (i.e., to classify a virus genome sequence as either severe or mild). Additionally, from the classification pipeline, important features are identified as Sites of Interest (SoIs) in the virus genome sequences for our downstream analysis.
2. Several CNN-RNN based models are used to predict mutations at specific Sites of Interest (SoIs) of the SARS-CoV-2 genome sequence followed by further analyses of the same on several South-Asian countries.
3. Overall, we present an analysis pipeline, shown in Figure 1, that can be further utilized as well as extended and revised (a) to analyse its virulence (e.g., with respect to the number of deaths its predecessors have caused in their respective countries) and (b) to analyse the mutation at specific important sites of the viral genome.

**Figure 1:**
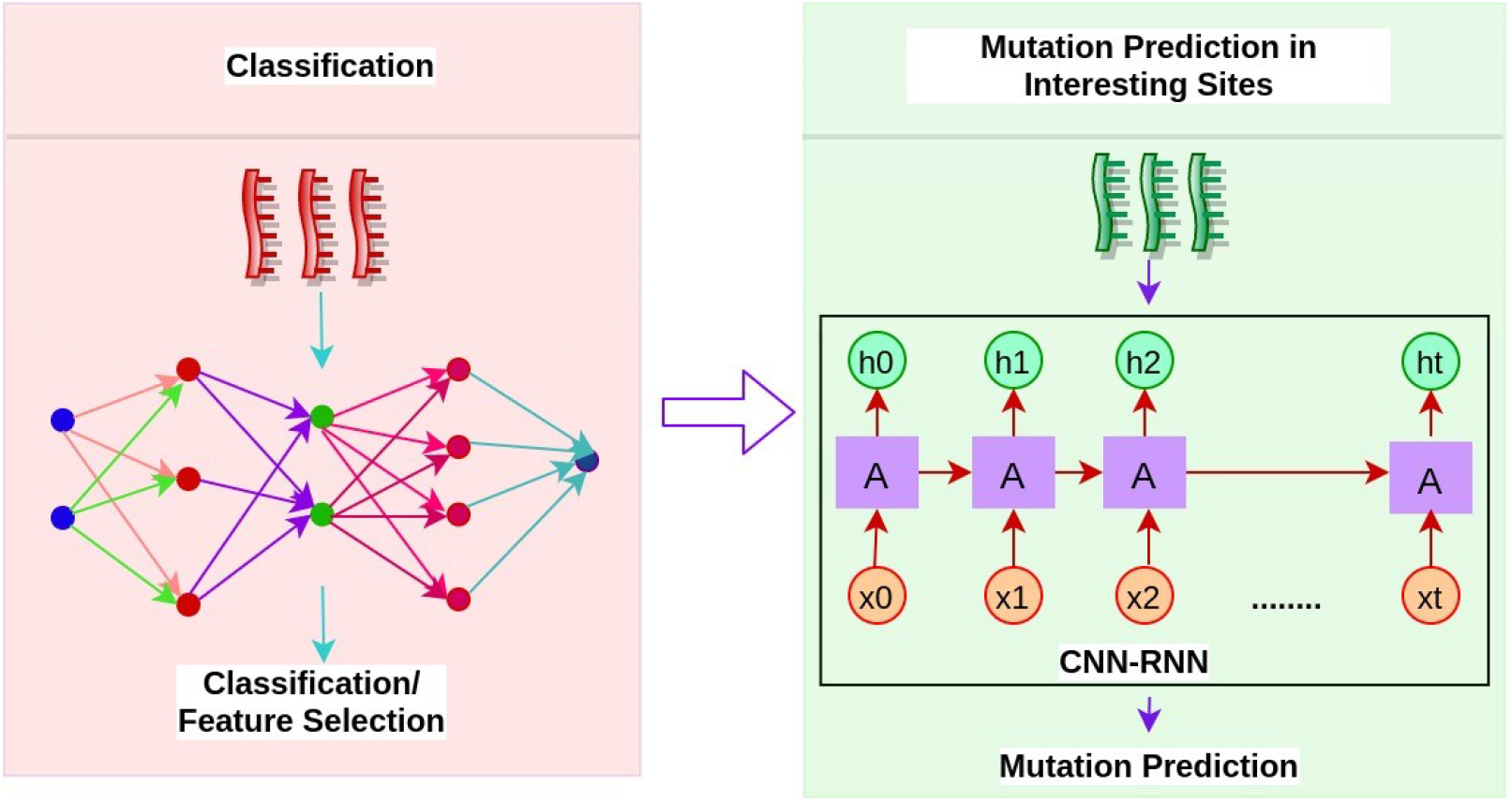
The whole analysis pipeline consisted of two phases. In the first phase, we have employed state-of-the-art classification algorithms, leveraging both traditional and deep learning pipelines to learn to discriminate the viral genome sequences of many countries as either mild or severe. We also identify the features that contributed the most as the discriminant factor in the classification pipeline. Then, we use the identified features from the previous stare to predict the mutation of the interesting sites in the viral genome sequence using a deep learning model.

## 2 Methodology

### 2.1 Data Collection and Preprocessing

We have collected 10179 hCov genome sequences upto the date 24 April, 2020 (cut-off date) from the GISAID initiative dataset [5]. These are high quality complete viral genome sequences submitted by the scientists and scientific institutes of individual countries. We will refer to the above dataset as Dataset A. Subsequently, to analyze and test our classification and mutation pipeline, we have collected all published genome sequences of several South-Asian countries upto 27 June, 2020 (Dataset B).

We also have collected country-wise death statistics (upto the cut-off date) from the official site of the World Health Organization [6]. The label is assigned based on a threshold of deaths which is the estimated median of the number of deaths in the data points. Any genome sequence of a country having deaths below (above) the threshold is considered a mild (severe) genome sequence, i.e., assigned a label 0 (1). A sample labelling is shown in the supplementary Table 1. Informatively, we have also considered some other metrics for labeling purposes albeit with unsatisfactory output (please see section 1.2 of the supplementary file for details). We have divided Dataset A into training and testing subsets in 80/20 ratio with a balanced number of data points per class for the traditional machine learning pipeline and for the deep learning classification routine, we have created the subsets training/validation/testing in 68/12/20 ratio.

### 2.2 Classification Models

#### 2.2.1 Traditional Machine Learning Pipeline

For traditional machine learning, we use a pipeline similar to [7] (See Figure 2). We extract three types of features from the genomic sequence of SARS-CoV-2. Inspired by the recent works [7–10] that focus only on sequences, we also extract only sequence based features. These features are: position independent features, *n*-gapped dinucleotides and position specific features (see details in Section 2 of supplementary file). We use the gini value [11] of the Extremely Randomized Tree (Extra Tree) classifier [12] to rank the features. Subsequently, only the features with gini value greater than the mean of the gini values are selected for training a LightGBM classifier model [13] (with default parameters) and 10-fold cross validation is performed. LightGBM is a highly efficient and fast gradient boosting framework which uses tree based algorithms.

**Figure 2:**
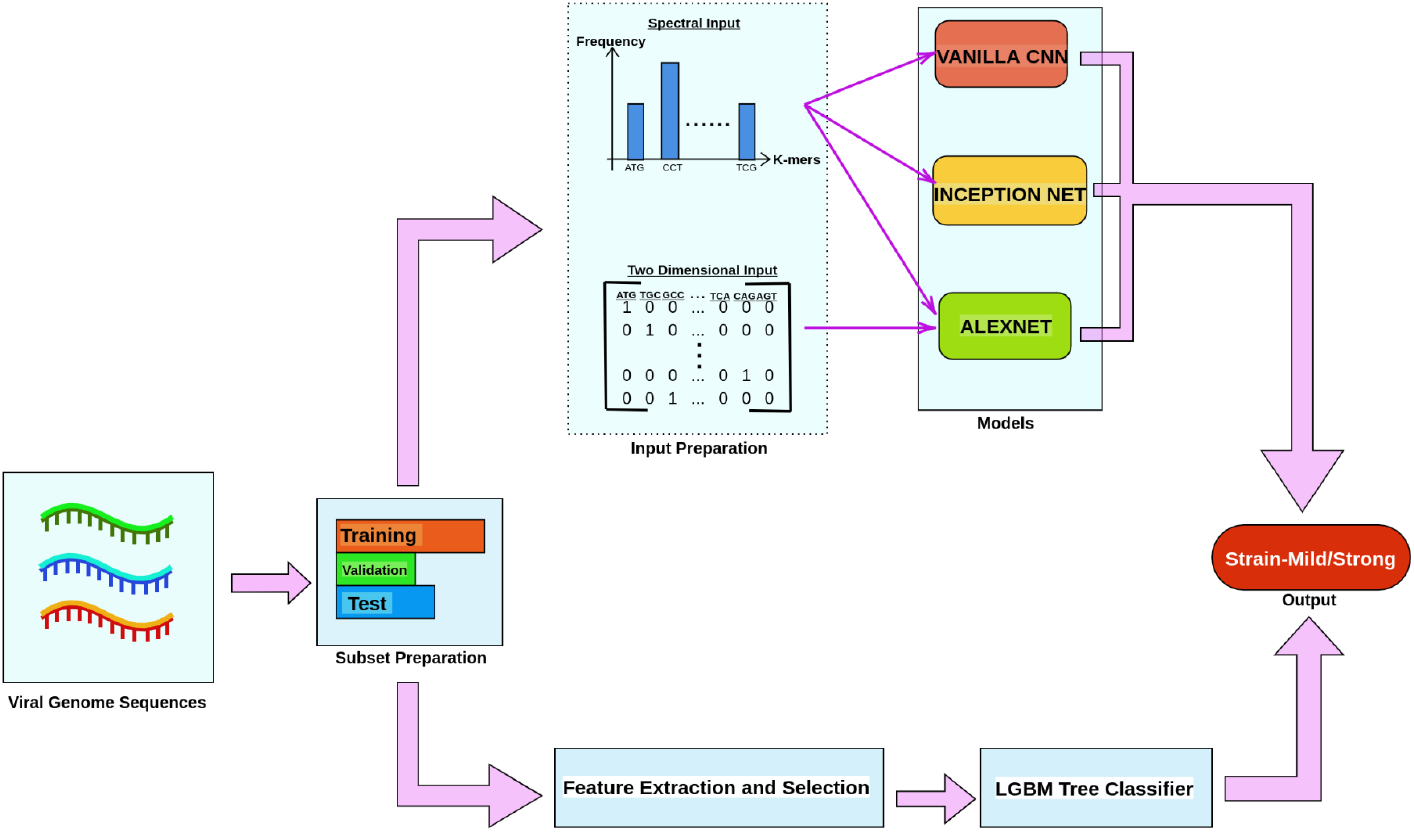
This figure illustrates our classification pipelines. We have leveraged the potential of both traditional and deep learning pipelines and in addition to doing an interesting classification task (mild vs. severe) we aim to identify, more importantly, the (sequence based) features and hence the corresponding sites of the viral genome that are performing as the most discriminant feature with respect to classification.

#### 2.2.2 Important feature identification

We use SHAP values and Univariate feature selection to compare the importance of the features. SHAP (SHapley Additive exPlanations) is a game theoretic approach which is used to explain the output of a model [14]. Univariate feature selection works by selecting the best features based on univariate statistical tests [15]. We use SelectKBest univariate feature selection to get the top K highest scoring features according to ANOVA f_classif feature scoring [16] function.

#### 2.2.3 Deep Learning Models

We leverage the power of 3 different deep learning (DL) classification models, namely, vanilla CNN [17], AlexNet [18] and InceptionNet [19]. We transform the raw viral genome sequences into two different representations, namely, K-mers spectral representation [17] and one hot vectorization [20] to feed those into the Deep Learning networks in a seamless manner (Figure 2). Details of these representations are given in Section 4.2 of the Supplementary File. For K-mers spectral representation we have experimented with different values of K (K = 3,5,7 for Vanila CNN and K = 3 & 5 only for the other models due to resource limitation). For one hot vectorization, we have trained InceptionNet for 150 epochs for both 3-and 5-mers and trained AlexNet for 135, 100 and 100 epochs for 3-,4-and 5-mers respectively.

### 2.3 Identifying the Representative Viral Genome Sequence

We aim to identify the representative viral genome sequences for the mutation prediction pipeline. To do that we have used an alignment-free genome sequence comparison method as proposed in [21], which is briefly described below and shown in Figure 3. Notably, we do not consider any alignment-based method since it is not computationally feasible for us to align thousands of viral sequences for analysis and clustering purposes [22].

**Figure 3:**
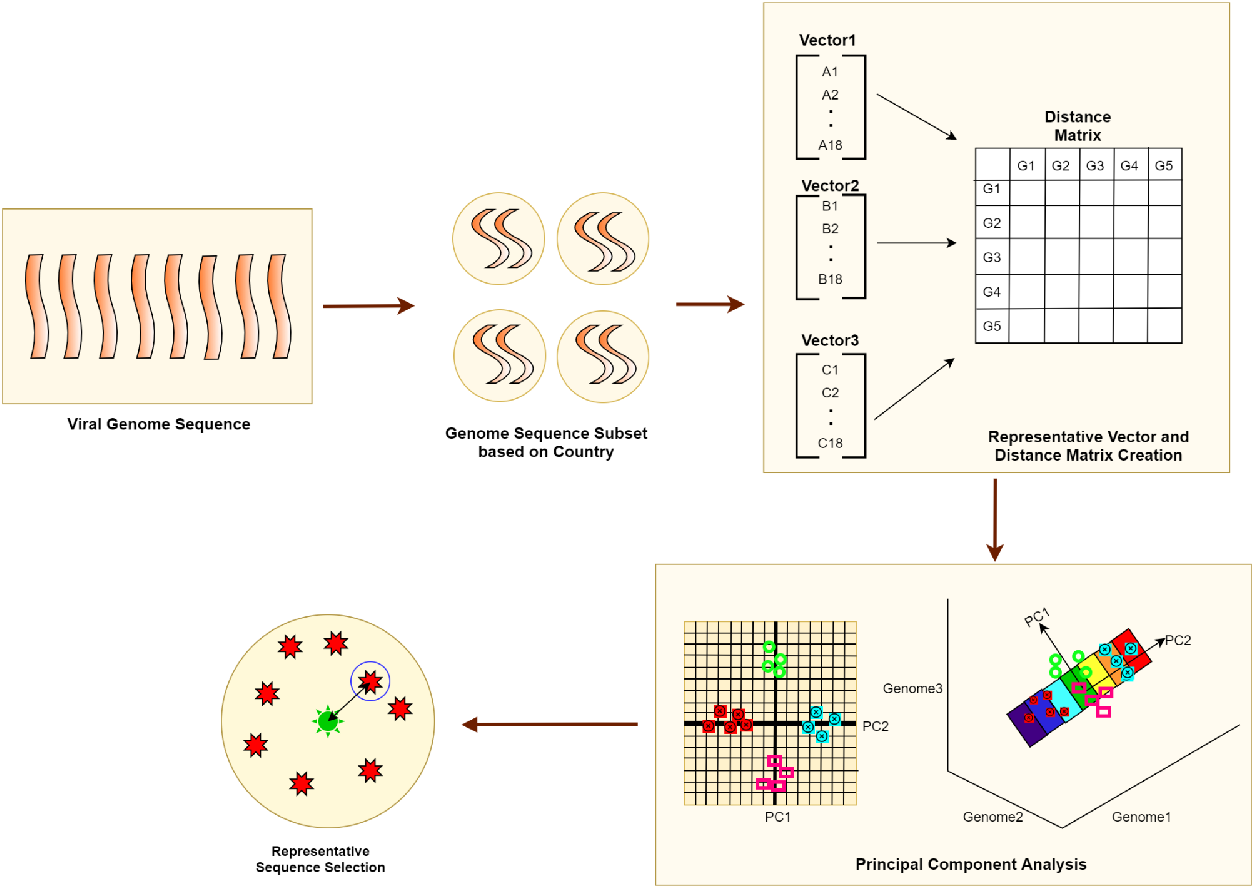
The Viral Genome Sequences were divided into subsets of sequences based on country. For each subset, each Viral Genome Sequence is converted into a vector representation and pairwise euclidean distance was calculated among the vectors to create the distance matrix. As the matrix is very high-dimensional, we used principal component analysis to find the principal component matrix from the distance matrix. Representative sequences were identified through K-means clustering on the PCA Matrix.

At first the sequence set is divided into subsets of sequences based on the location. All sequences are converted into representative IR^18^ vectors. Pairwise distance among vectors derived from the fast vector method [21] are computed using Euclidean distance. Due to the high dimensionality of the resulting distance matrix, we resort to the Principal Component Analysis (PCA) technique [23] to reduce the dimension of the matrix. Subsequently, we use K-means clustering [24] to identify the corresponding cluster centers. For the K-means clustering algorithm, we have used the implementation of [16] and used the default parameters except for the number of clusters which were set to 1 for determining the cluster center for each of the subsets. For each location-based cluster, the representative sequence (i.e., the “centroid” of the cluster) is then identified and used in the subsequent step of the pipeline.

### 2.4 Mutation Prediction

We design a pipeline shown in Figure 4 to predict mutation on the sites of interest (as identified through our classification pipeline) in the SARS-CoV-2 genome. We follow a similar protocol followed by [25] and adopt it to fit our setting as follows. We divide all the available countries and the states of the USA into different time-steps by the date of the first reported incidence of SARS-CoV-2 infected patients of that location. Thus, every resulting time-step represents a date (*T_k_* for Cluster *k*) and contains the clusters of genome sequences of the respective countries/states. Then the time series samples are generated by concatenating sites from different time-steps one-by-one that represent the evolutionary path of the SARS-CoV-2 viral genome sequence. For example, *T*_1_ is the very first date when the virus is discovered in China. So, time-step 1 contains only one country, China. Likewise, time-step 2 contains clusters for those countries where the virus is discovered on date *T*_2_ and so on. We generate 300000 time series sequences by concatenating genome sites from *T*_1_,*T*_2_,….,*T_n_* (in our case, *n =* 40), divide the samples in 68/12/20 ratio and then feed the training samples to the model that consists of a convolutional one dimensional layer and a recurrent neural network layer [26]. In our experiments, we have used both pure LSTM and bidirectional LSTM as our RNN layer (see Section 3.3 of supplementary file). The model has a dense layer of 4 neurons in the end which predicts the probability of the next base pair of the next time-step. So, in a nutshell, the model takes concatenated genome sequences from *T*_1_,*T*_2_,·,*T_n−_*_1_ as input and predicts the mutation at time *T_n_*.

**Figure 4:**
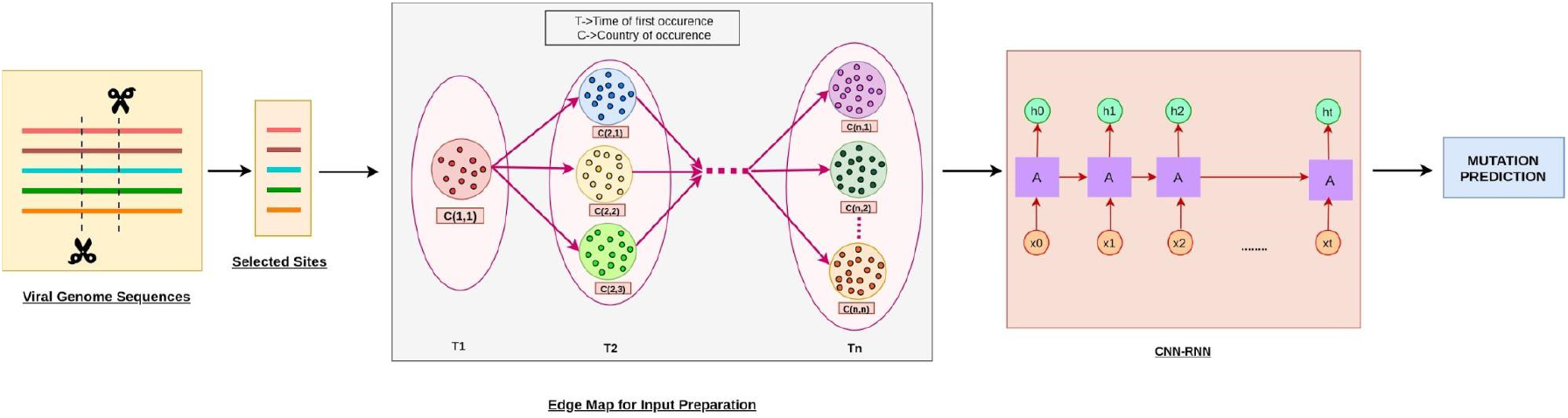
The interesting sites were selected from the viral genome sequences using the feature selection routine described in the classification methods. The sites were then divided by geographical locations and clustered by the time of the first occurrence at that respective region. Time-series sequences were created by concatenating random genome sequences from the closest sub-clusters and trained in an CNN-RNN network for predicting mutation in the sites of the final time-step.

We further use our mutation prediction pipeline to identify and analyze possible parents of a mutated genome sequence. For this particular analysis, we trained the models specifically for some South-Asian countries, namely, Bangladesh, India and Pakistan. We only used the best performing model for this analysis and generated five time series samples. At the time of generating these samples, the country/location having the minimal euclidean distance with the country/location of the next time-step was taken for each time-step.

### 2.5 Coding and Experimental Environment

We have implemented our experiments mostly in python. We have used scikit-learn library [16] for clustering and plotting the graphs. For deep learning models, scikit-learn, TensorFlow and keras neural network libraries are used and for LightGBM classifier, python LightGBM framework has been used. The experiments have been conducted in the following machines:

- Experiments involving the deep learning pipelines (i.e., both classification and mutation prediction) have been conducted in the work-stations of Galileo Cloud Computing Platform [27] and the default GPU provided by the Google Colaboratory Cloud Computing Platform [28].
- The LightGBM classifier model was trained in a machine with Intel Core i5-4010U CPU @ 1.70GHz x 4, Windows 10 OS and 16 GB RAM.

All the codes and data (except for the Genome Sequences) of our pipeline can be found at the following link: https://github.com/pythonLoader/Analyzing-hCov-Genome-Sequence. The Genome Sequence data have been extracted from and are publicly available at GISAID [5].

## 3 Results

### 3.1 Mild/Severe genome sequence Classification

All our classifiers are trained to learn whether a given genome sequence is mild or severe. The classification accuracy of the LightGBM classifier (97%) is superior to that of the deep learning classifiers (84-89%), which, while is somewhat surprising, is inline with the recent findings of [7]. It should be noted that LightGBM had produced better results in significantly less time than deep learning models for this dataset. The results of the classifier models are shown in Figure 5.

**Figure 5:**
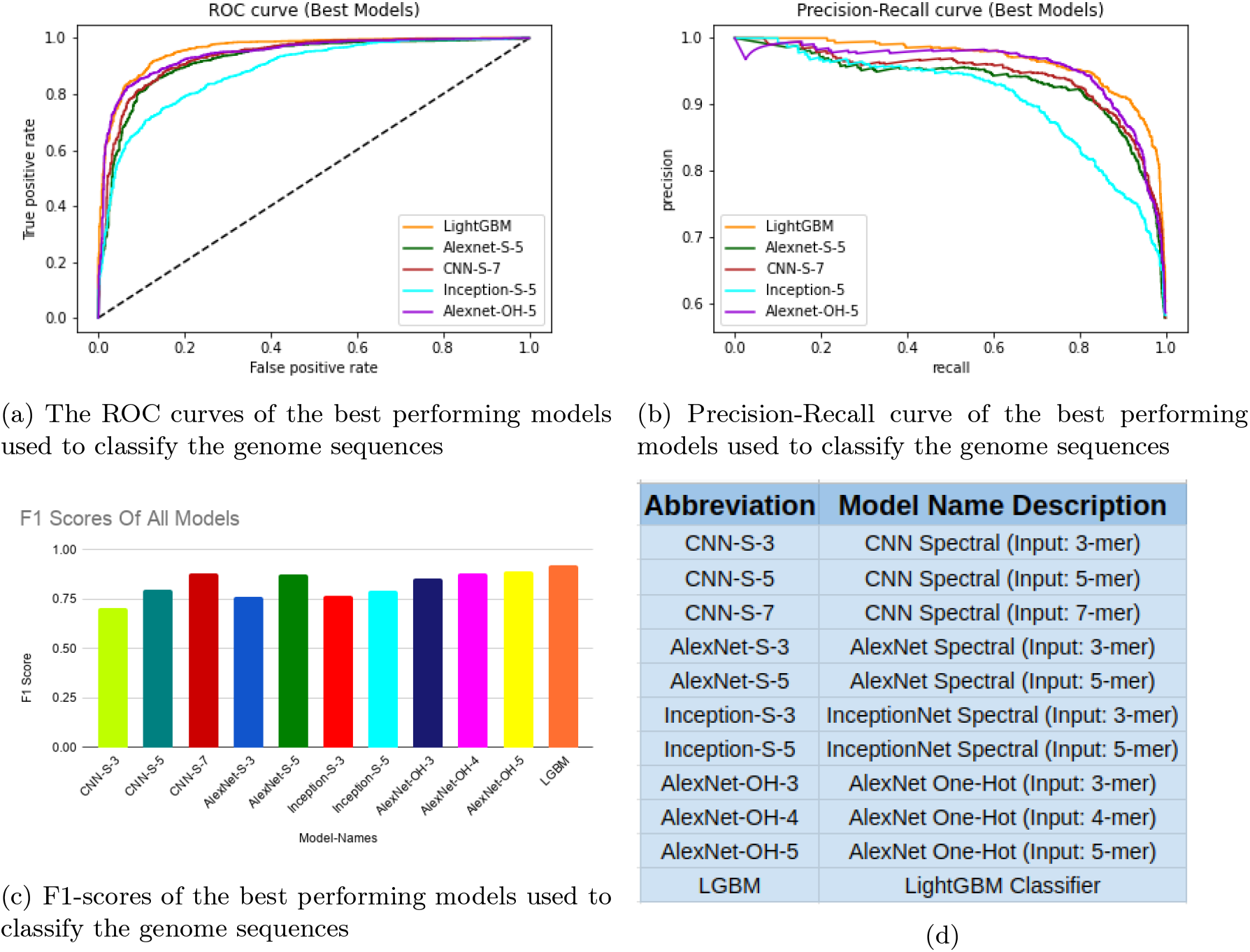
The Receiver Operating Characteristic Curves (ROC) show the diagnostic capability of the ML and the DL classifiers used in our experiments. Please refer to Table 5 and Figure 3 for detailed results.

Quantitative results aside, we also have applied our classifiers on the sequences that have been deposited at GISAID after the cut-off date (i.e., April 24, 2020). Since the cut-off date, the country wise death statistics [6] has certainly changed significantly and this has pushed a few countries, particularly from Asian regions and several states of the United States of America to transition from mild to severe states (based on our predefined threshold). Interestingly, our classifiers have been able to predict the severity of the new genome sequences submitted from these countries/states correctly. Table 6 in the supplementary file shows a snapshot of a few such countries/states with the relevant information. Furthermore, 957 new genome sequences from India, 151 genome sequences from Bangladesh and 3 genome sequences of Pakistan that were collected well after the cut-off date upto 27 June, 2020 were classified as severe sequences with 100% accuracy by the LGBM classifier.

### 3.2 Sites of Interest (SoI)

We preliminarily identify the top 10 features of SHAP and SelectKBest feature selection (with *k =* 10). From these features, as SoIs, we have selected the features that are also biologically significant, i.e., cover different significant gene expression regions (Figure 6). In particular, we have selected the position specific features pos_8445_8449, pos_19610_19614, pos_24065_24069 and pos_23825_23829 as the SoIs for the mutation prediction analyses down the pipeline. Here, pos_X_Y indicates the site from Positions X to Y of the virus genome sequences. The reason for selecting these features as SoIs are outlined below.

According to gene expression studies [29,30], our SoIs, namely, pos_8445_8449 and pos_19610 _19614 encode to two Non-structural Proteins, Nsp3 and Nsp11, respectively. Also, our other two SoIs, namely, pos_24065_24069 and pos_23825_23829 correspond to the Spike Protein of SARS-CoV-2. Nsp3 binds to viral RNA, nucleocapsid protein, as well as other viral proteins, and participates in polyprotein processing. It is an essential component of the replication/transcription complex [31]. So, the mutation in this protein is expected to affect the replication process of the SARS-CoV-2 in host bodies. On the other hand, the spike protein sticks out from the envelope of the virion and plays a pivotal role in the receptor host selectivity and cellular attachment. According to Wan et al. there exists strong scientific evidence that SARS and SARS-CoV-2 spike proteins interact with angiotensin-converting enzyme 2 (ACE2) [32]. The mutation on this protein is expected to have a significant impact on the human to human transmission [33]. Therefore, it is certainly interesting and useful to predict the mutation of such SoIs.

**Figure 6:**
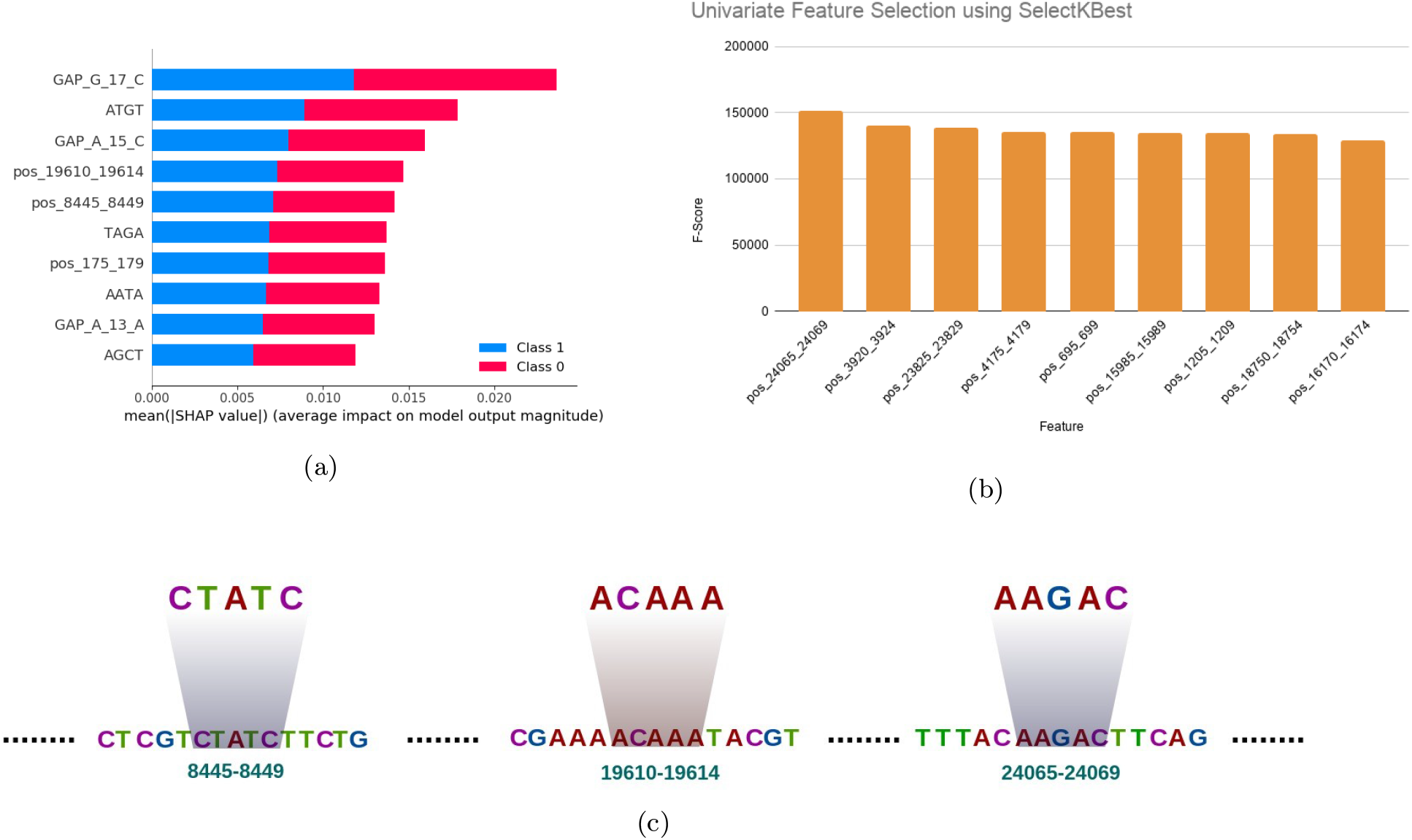
Top 10 features based on SHAP values(a) and SelectKBest(b) and identifying the position specific feature(s) from the genome sequence as site(s) of interest(c). These SOIs will be used for mutation prediction down the pipeline.

### 3.3 Mutation Prediction Results

CNN-LSTM and CNN-bidirectional LSTM performed similarly for different SoIs of the genome. Except for a few positions (i.e., mostly the first couple of positions) in the SoIs, the performance of the mutation prediction pipelines are quite promising. In particular, for the positions 19610 to 19614, both the pipeline performed fairly keeping the accuracy above 0.7 (the best accuracy of 79.89% was achieved by both of them for position 19611). For the positions 8447, 8448 and 8449, CNN-bidirectional LSTM performed slightly better achieving 87.35%, 89.95% and 92.69% accuracy respectively. For the positions 24067, 24068 and 24069, both the pipeline performed identically achieving 92.33%, 87.43% and 92.48% accuracy respectively. For the positions 23827 and 23828, CNN-bidirectional LSTM performed slightly better achieving 84.87% and 95% accuracy respectively and for position 23829, both performed identically with accuracy 89.75%. For detailed results (please check Table 7 of the supplementary material).

### 3.3.1 Improving the performance on the first few sites

As has been mentioned above, the performance of the mutation prediction pipeline is not up to the mark for the first couple of positions in the SoIs in most cases whereas the performance is excellent for the third position onward. To improve the mutation prediction of the first two positions of the relevant sites of interest (SoIs), we further trained the best model (CNN-Bidirectional LSTM) for the SoI starting from the preceding three positions (to the first two positions thereof) thereby feeding more information to the architecture with respect to the first two positions. Consequently, the accuracy of the prediction improved by a substantial margin (please see Table 10 in the supplementary file).

### 3.3.2 Analyzing Parent genome sequences

For the analysis involving only Bangladesh, we used the CNN-bidirectional LSTM model (as this performed slightly better between the two) and achieved almost 100% accuracy. Then we analyzed the ancestors in the time series test samples and noticed that some of the states of the USA are present in these samples. These states are California, Massachusetts, Texas, New Jersey and Maryland. For India and Pakistan, we got similar results for some sites but for other sites, accuracy was not as high as that of Bangladesh (Check Table 8 of the supplementary file for details).

### 3.3.3 Mutation Prediction on new SARS-CoV-2 genome sequence

We analyzed the genome sequences of Bangladesh collected after the cut-off date. Our deep learning pipeline was able to predict the mutations in the SoI of the spike protein region (pos_24065_24069) with 80-86% accuracy. (Please refer to Table 9 of the supplementary file).

## 4 Discussions

Our analysis pipeline revealed some interesting and insightful findings as discussed below. A genome sequence (EPI_ISL_435050) was collected on April 13, 2020 (before our cut-off date) from a patient in Ahmedabad, Gujrat, India. It was predicted to be a severe genome sequence (with low confidence) even though we trained the classifier to consider the Indian sequences as mild. We considered another genome sequence (EPI_ISL_437447) which was collected from another patient from the same place in India on April 26, 2020 (after our cut-off date) and predicted the severity thereof. The classifiers declared this isolate to be severe with very high confidence (about 98%). This strongly suggests that there were some mutations that turned the Indian sequences from mild or less severe to severe or highly severe, respectively.

Our mutation prediction pipeline originally performed better for the third, fourth and fifth positions for most SoIs (Please check Table 8 of the supplementary file) and mostly failed for the first two positions. Through some analysis, we conjectured that for the first couple of positions the models are not getting enough information to work on. Subsequently, we verified our conjecture through further experiments. Thus it can be concluded that given more information at the time of prediction such as providing the first two positions at the time of predicting the third, the RNN-LSTM models can predict more accurately.

Also, we conducted an analysis to predict possible parents of the (mutated) virus genome sequences of the South Asian Region (Bangladesh, India and Pakistan). Our mutation prediction pipeline suggested that the genome sequences of some states of the USA, namely, California, Massachusetts, Texas, New Jersey and Maryland could be the parents/ancestors of these South Asian genome sequences. This could explain the COVID-19 surge (27 cases per 1,00,000 people) in South Asian countries during the middle part (June-July) of the year 2020 [34].

Our current mutation prediction pipeline is only based on some specific sites of interest. However, it promises to be extremely useful in predicting the mutations in the viral strain. In fact, it can potentially be used to predict mutated strains considering that each position has the possibility to mutate. However, this would require huge computational power which we lack. A better approach of course would be to check only biologically meaningful sites of interests for possible mutation. To this end, we analyzed the new Bangladeshi (BD) genome sequences and found that our mutation prediction pipeline predicted the SoI (pos_24065_24069) to be AACAA which matched with 80% of the test BD genome sequences.

Finally, from a computational perspective a brief comment on the lower performance of the deep learning pipelines (against the traditional machine learning ones) is in order. Perhaps, the true potential of the deep learning pipelines will unfold if and when more data will be fed thereto which we could not manage to do due to our computational resource constraints.

## 5 Limitations of the Study

The study has several potential limitations. Our estimate ignores the environmental and geographical aspect of the place where the virus is spreading. The mutation prediction can only be carried out in particular sites of interest as whole sequence prediction has a serious technological bottleneck. Furthermore, we could not increase our sample space to improve the accuracy of our models as a result of limited computational resources. Finally, we did not have unbounded access to some health related information such as recovery rate, the health infrastructure of a country and the governmental strategy to fight this pandemic to label the genome sequences properly.

## 6 Future Works

We believe that out analysis pipeline will spark more research endeavours using and improving both our classification and mutation pipelines. As an immediate extesnion the following works could be condiered.

1. Through a detailed latent factor analysis, other alternate and improved labeling scheme with all the new genome sequences present in GISAID archive should be studied. This will improve the overall classification pipeline and perhaps identify more SoIs with biological significance.
2. New mutation prediction architecture can be tried out to improve the mutation prediction accuracy, e.g., experimenting with transition matrix concepts used in Route Prediction or Credit Rating Analysis and applying such concepts in this domain.
3. Another immediate task could be to feed more data to our deep learning classification pipelines and unfold the true potential thereof which we could not do due to our computational constraints.

