## Supplementary Material for "Analyzing hCov Genome Sequences: Predicting Virulence and Mutation"

#### 1 Input Preprocessing and Labeling

##### 1.1 Primary Labeling

For mild vs. severe classification, primarily, the label was assigned based on a threshold of deaths, i.e., the estimated median of the number of deaths in the data points. For our data points this threshold was 2000. Any genome sequence of a country having deaths below (above) the threshold were considered a mild (severe) strain, i.e., assigned a label 0 (1). A sample labeling is shown in Table 1.

##### 1.2 Alternative Labeling

Informatively, we have also considered some other metrics for labeling purposes albeit with unsatisfactory output such as death rate based on recovery that is case fatality rate recovery case fatality rate recovery or (CFR_Recovery), death rate based on number of confirmed cases (CFR_Infection) and labeling based on the weighted health infrastructure information (CFR_Infrastructure). The composite index from [1] is considered as the weighted health infrastructure information. The composite index is calculated as the weighted average of five component goals namely health, health inequality, responsiveness-level, responsiveness-distribution, and fair-financing. These metrics are given below:

$$CFR_{Recovery} = \frac{Number\ of\ deaths}{Number\ of\ deaths + Number\ of\ recoveries}$$

$$CFR_{Infection} = \frac{Number\ of\ deaths}{Number\ of\ deaths + Number\ of\ infections}$$

$$CFR_{Infrastructure} = CFR_{Infection} * x + (1 - x) * composite\ index$$

Here, $x$ is calculated from a linear regression model. First we have selected 15 countries for which label (CFR_Infrastructure) could be assumed with confidence. Solving the equation for CFR_Infrastructure, $x$ was calculated. Using these 15 data points we have developed a linear regression model to predict the value of $x$ when CFR_Infection and composite index is available as input.

The resulting labels were counter-intuitive. In case of CFR_Recovery, some strains (e.g., Chinese, Iranian) which should be severe became mild and other metrics did not produce satisfactory results. Moreover, the difference between the median and other labeling techniques is menial, e.g. ( 16% with CFR_Recovery, 11% with both CFR_Infection and CFR_Infrastructure). This is shown in table 2

Table 1: Labeling genome sequences based on the number of deaths. The threshold (estimated median of the data points) was 2000, so any genome sequences of a country having deaths below (above) the threshold were considered a mild (severe) strain.

| **Countries** | **Deaths** | **Label** | **Countries** | **Deaths** | **Label** |
|---|---|---|---|---|---|
| Algeria | 407 | 0 | Hungary | 250 | 0 |
| Democratic Republic of the Congo | 25 | 0 | Iceland | 10 | 0 |
| Gambia | 1 | 0 | Ireland | 794 | 0 |
| Ghana | 9 | 0 | Italy | 25549 | 1 |
| Nigeria | 31 | 0 | Latvia | 11 | 0 |
| Senegal | 6 | 0 | Lithuania | 40 | 0 |
| South Africa | 75 | 0 | Luxembourg | 83 | 0 |
| Cambodia | 0 | 0 | Netherlands | 4177 | 1 |
| China | 4642 | 1 | Norway | 180 | 0 |
| Georgia | 5 | 0 | Poland | 454 | 0 |
| India | 718 | 0 | Portugal | 820 | 0 |
| Indonesia | 647 | 0 | Slovakia | 15 | 0 |
| Israel | 192 | 0 | Slovenia | 79 | 0 |
| Japan | 317 | 0 | Spain | 22157 | 1 |
| Kuwait | 14 | 0 | Sweden | 2021 | 1 |
| Malaysia | 95 | 0 | Switzerland | 1267 | 0 |
| Nepal | 0 | 0 | Turkey | 2491 | 1 |
| Pakistan | 237 | 0 | Canada | 2028 | 1 |
| Philippines | 462 | 0 | Mexico | 970 | 0 |
| Qatar | 10 | 0 | Australia | 76 | 0 |
| Saudi Arabia | 121 | 0 | New Zealand | 17 | 0 |
| Singapore | 12 | 0 | Argentina | 159 | 0 |
| Sri Lanka | 7 | 0 | Brazil | 2906 | 1 |
| Thailand | 50 | 0 | Chile | 168 | 0 |
| Panama | 144 | 0 | Colombia | 206 | 0 |
| Austria | 508 | 0 | Ecuador | 560 | 0 |
| Belarus | 60 | 0 | Peru | 530 | 0 |
| Belgium | 6490 | 1 | Vietnam | 0 | 0 |
| Denmark | 394 | 0 | Iran | 5481 | 1 |
| Estonia | 45 | 0 | South Korea | 240 | 0 |
| Finland | 172 | 0 | USA | 42311 | 1 |
| France | 21823 | 1 | United Kingdom | 18738 | 1 |
| Germany | 5321 | 1 | Russia | 615 | 0 |
| Greece | 125 | 0 | | | |

Table 2: Alternate labeling of the input set and the discrepancy with the original labeling, three other labeling were used which are death rate based on recovery, and death rate based on number of deaths by number of confirmed cases, labeling based on the weighted health infrastructure information and death rate. The entries marked in green represent that the alternate labeling indicates severe whereas the original label indicates mild and the red ones are for vice-versa.

| **Countries** | **Indicator_death (Originally used)** | **Indicator_CFR_ Recovery** | **Indicator_CFR_ Infection** | **Indicator_Infrastructure** |
|---|---|---|---|---|
| Algeria | 0 | 0 | 1 | 1 |
| Congo | 0 | 1 | 1 | 0 |
| Ghana | 0 | 0 | 0 | 0 |
| Nigeria | 0 | 0 | 0 | 0 |
| Senegal | 0 | 0 | 0 | 0 |
| South Africa | 0 | 0 | 0 | 0 |
| Cambodia | 0 | 0 | 0 | 0 |
| China | 1 | 0 | 1 | 1 |
| Georgia | 0 | 0 | 0 | 0 |
| India | 0 | 0 | 0 | 0 |
| Israel | 0 | 0 | 0 | 0 |
| Japan | 0 | 0 | 0 | 0 |
| Kuwait | 0 | 0 | 0 | 0 |
| Malaysia | 0 | 0 | 0 | 0 |
| Nepal | 0 | 0 | 0 | 0 |
| Pakistan | 0 | 0 | 0 | 0 |
| Qatar | 0 | 0 | 0 | 0 |
| Saudi Arabia | 0 | 0 | 0 | 0 |
| Singapore | 0 | 0 | 0 | 0 |
| Thailand | 0 | 0 | 0 | 0 |
| Panama | 0 | 1 | 0 | 0 |
| Austria | 0 | 0 | 0 | 0 |
| Belarus | 0 | 0 | 0 | 0 |
| Belgium | 1 | 1 | 1 | 1 |
| Denmark | 0 | 0 | 1 | 1 |
| Estonia | 0 | 0 | 0 | 0 |
| Finland | 0 | 0 | 0 | 0 |
| France | 1 | 1 | 1 | 1 |
| Germany | 1 | 0 | 0 | 0 |
| Greece | 0 | 0 | 1 | 1 |
| Hungary | 0 | 1 | 1 | 1 |
| Iceland | 0 | 0 | 0 | 0 |
| Ireland | 0 | 0 | 0 | 0 |
| Italy | 1 | 1 | 1 | 1 |
| Latvia | 0 | 0 | 0 | 0 |
| Lithuania | 0 | 0 | 0 | 0 |

| | | | | |
|---|---|---|---|---|
| Luxembourg | 0 | 0 | 0 | 0 |
| Netherlands | 1 | 1 | 1 | 1 |
| Norway | 0 | 1 | 0 | 0 |
| Poland | 0 | 0 | 0 | 0 |
| Portugal | 0 | 1 | 0 | 0 |
| Slovakia | 0 | 0 | 0 | 0 |
| Slovenia | 0 | 0 | 1 | 1 |
| Spain | 1 | 0 | 1 | 1 |
| Sweden | 1 | 1 | 1 | 1 |
| Switzerland | 0 | 0 | 0 | 0 |
| Turkey | 1 | 0 | 0 | 0 |
| Canada | 1 | 0 | 0 | 0 |
| Mexico | 0 | 0 | 1 | 1 |
| Australia | 0 | 0 | 0 | 0 |
| New Zealand | 0 | 0 | 0 | 0 |
| Argentina | 0 | 0 | 0 | 1 |
| Brazil | 1 | 0 | 1 | 1 |
| Chile | 0 | 0 | 0 | 0 |
| Colombia | 0 | 0 | 0 | 0 |
| Ecuador | 0 | 0 | 0 | 0 |
| Peru | 0 | 0 | 0 | 0 |
| Vietnam | 0 | 0 | 0 | 0 |
| Iran | 1 | 0 | 1 | 1 |
| South Korea | 0 | 0 | 0 | 0 |
| USA | 1 | 1 | 1 | 1 |
| United Kingdom | 1 | 1 | 1 | 1 |
| Russia | 0 | 0 | 0 | 0 |

### 2 Feature Extraction and Selection

#### 2.1 Feature Extraction

##### 2.1.1 Position Independent Features

After the preprocessing steps the sequences contain 4 nucleotide symbols (A, C, T and G). With these four nucleotides different k-mer combinations are possible. For 2-mer, such combinations will be AC, AT, AG etc. In case of position independent features we counted the number of occurrences of a specific k-mer in the whole genome sequence. In this experiment we had extracted such features for up to 4-mers.

##### 2.1.2 n-Gapped Dinucleotides

In these types of features, we counted the occurrences of two nucleotides at a certain distance. For example, 2-gapped AC would count the occurrence of A and C two positions apart in the sequence.

For our experiment we considered the value of n to be from 1 to 30. These types of features were used in [2–4].

###### 2.1.3 Position Specific Features

In the previous studies [2, 3] these were binary features. They represented whether a k-mer exists in a specific position or not. For example if we consider 1-mer, for position 1 there will be four features for A, C, T and G. Thus a lot of features are generated which is feasible for small sequences like sgRNA [2, 5]. We note that for finding the position specific features, we had two major issues as described below:

1. The sequences in this dataset had variable length, ranging from 29201 to 30129 which might have caused a variable number of features for each sample.

2. The second problem was the sheer numbers of features. Even if we considered 1-mer features only, the number of features would have been more than 100000.

We solved the first problem by assuming that all the sequences had 29000 nucleotides. Those with more nucleotides were stripped and those with less had empty substring in the padded positions. The second problem was solved by partitioning the whole genome sequence into disjoint subsequences and one feature value was calculated for each subsequence. In our experiment the subsequences were 5-mers. Thus we reduced the number of these types of features to 6000. A 5-mer subsequence can have 1024 unique combinations. For each subsequence we assigned a number from 1 to 1024 depending on the combinations. If the subsequence was empty (for those sequences with length less than 30000) 0 was assigned as feature value.

These problems could have been avoided if we had all the genome sequences perfectly aligned with each other.

##### 2.2 Feature Selection

We ranked the features using the Extremely Randomized Tree (Extra Tree) classifier. The ranking criterion was gini value. Feature selection was performed to reduce the number of features. Only the features with gini value greater than the mean of the gini values were selected for training the model.

#### 3 Mutation prediction

##### 3.1 Input Preparation

5 genome sequences closer to the cluster centre was taken and it's sites of interest was extracted. Next according to the temporal date based stages, the sites were appended to form the mutation sequences which were used downstream.

Table 3: Date of first occurrence of the virus of a country at that date. The genome sequences of the countries were taken in country-specific clusters for every date.

| Date of First Occurence | Countries/ US States at that time stamp |
|---|---|
| 2019-11-17 | China |
| 2020-01-13 | Nepal, Thailand |
| 2020-01-16 | Japan |
| 2020-01-20 | South Korea |
| 2020-01-21 | Washington, Taiwan |
| 2020-01-23 | Singapore, Vietnam, Hong Kong |
| 2020-01-24 | France, Illinois |
| 2020-01-25 | Malaysia, Australia, California |
| 2020-01-26 | Arizona |
| 2020-01-27 | Cambodia, Sri Lanka, Germany, Canada |
| 2020-01-29 | Pakistan, Finland |
| 2020-01-30 | India, Philippines |
| 2020-01-31 | Italy, Spain, Sweden, United Kingdom, Russia |
| 2020-02-01 | Massachusetts |
| 2020-02-03 | Senegal, Indonesia, Saudi Arabia, Latvia, Portugal |
| 2020-02-04 | Belgium |
| 2020-02-05 | Wisconsin |
| 2020-02-12 | Texas |
| 2020-02-17 | Nebraska |
| 2020-02-19 | Iran |
| 2020-02-21 | Israel |
| 2020-02-24 | Kuwait |
| 2020-02-25 | Algeria, Austria, Switzerland, Brazil, Utah |
| 2020-02-26 | Georgia, Greece, Norway |
| 2020-02-27 | Nigeria, Qatar, Denmark, Estonia, Netherlands |
| 2020-02-28 | Belarus, Iceland, Lithuania, New Zealand, Oregon, Mexico |
| 2020-02-29 | Ireland, Luxembourg, Ecuador |
| 2020-03-01 | Czech Republic, Florida, New York, Rhode Island |
| 2020-03-02 | USA_Georgia, New Hampshire |
| 2020-03-03 | Argentina, Chile, North Carolina |
| 2020-03-04 | New Jersey, Hungary, Poland Slovenia |
| 2020-03-05 | Colorado, Maryland, Nevada, Tennessee, South Africa |
| 2020-03-06 | Hawaii, Indiana, Kentucky, Minnesota, Oklahoma, Pennsylvania, South Carolina, Slovakia, Colombia, Peru |
| 2020-03-07 | District of Columbia, Kansas, Missouri, Vermont, Virginia |
| 2020-03-08 | Connecticut, Iowa |
| 2020-03-09 | Louisiana, Ohio, Panama |
| 2020-03-10 | Democratic Republic of the Congo, Michigan, South Dakota |
| 2020-03-11 | Arkansas, Delaware, Mississippi, New Mexico, North Dakota, Wyoming, Turkey |
| 2020-03-12 | Ghana, Alaska, Maine |
| 2020-03-13 | Alabama, Idaho, Montana, Puerto Rico, Uruguay |

#### 3.2 Experimentation details

Every country wise cluster contains maximum 20 genome sequences.These are the closest 20 genomes to the cluster centers.

#### 3.3 Model Description

First, all nucleotides of the concatenated time series samples are converted into one-hot vectors. Then, the input is fed to a one dimensional convolutional layer with 1024 filters.This layer is followed by a max pooling and a dropout layer.To learn the time series samples better,we then added an RNN layer(in one model it is vanilla LSTM and in another, bidirectional LSTM layer is added) which is followed by another dropout layer. Finally the flattened output was fed to two dense layers with 1024 and 4 neurons each.

### 4 Classification Models

#### 4.1 LGBM Classifier Model Pipeline

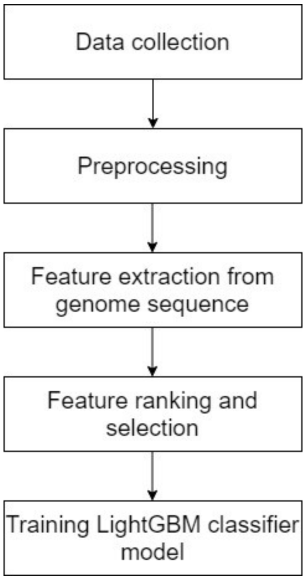


Figure 1: Steps for training LGBM Classifier Model

##### 4.1.1 LightGBM Classifier

Gradient boosting is a method which uses the ensemble of weak predictors. Generally these predictors are decision trees. In gradient boosting a differentiable loss function is optimized. LightGBM is a framework for gradient boosting. It is very fast compared to other gradient boosting frameworks like XGBoost and uses less memory. For this experiment we have used the default parameters of LightGBM classifier.

##### 4.1.2 Extremely Randomized Trees (Extra Trees)

It is an ensemble learning method like random forest, but it is faster. The main difference is Extra Trees algorithm selects totally random value for split whereas random forest selects locally optimal split. We have used Extra Trees for ranking features and selecting a subset of them to reduce the dimension of the feature matrix. In the experiment we have used 'gini score' for feature ranking criterion.

##### 4.1.3 Cross Validation

We have performed 10 fold cross validation on the dataset using this pipeline. Each fold contained 20% samples as the test set. The data was shuffled before split and the splits were stratified by data labels. Result of cross validation is shown in Table 4.

Table 4: Results of the 10-fold cross validation is shown in the table. For each fold accuracy, precision, recall, F1 score and AUROC have been mentioned.

| Fold | Accuracy | Precision | Recall | F1 Score | AUROC |
|---|---|---|---|---|---|
| 1 | 0.8969578018 | 0.905 | 0.9187817259 | 0.9118387909 | 0.9579360185 |
| 2 | **0.9076620825** | **0.9147157191** | 0.9271186441 | 0.9208754209 | **0.9683629019** |
| 3 | 0.884086444 | 0.8806451613 | 0.9254237288 | 0.9024793388 | 0.9511880247 |
| 4 | 0.8988212181 | 0.8959349593 | 0.9338983051 | 0.9145228216 | 0.9543065896 |
| 5 | 0.8998035363 | 0.8986928105 | 0.9322033898 | 0.9151414309 | 0.9596744812 |
| 6 | 0.9086444008 | 0.901453958 | **0.9457627119** | **0.9230769231** | 0.9615594804 |
| 7 | 0.8968565815 | 0.8917609047 | 0.9355932203 | 0.9131513648 | 0.9539204815 |
| 8 | 0.8899803536 | 0.8943894389 | 0.9186440678 | 0.9063545151 | 0.9498613971 |
| 9 | 0.8997050147 | 0.9053156146 | 0.9237288136 | 0.9144295302 | 0.9518219347 |
| 10 | 0.9046214356 | 0.9087893864 | 0.9288135593 | 0.9186923722 | 0.9621799706 |
| **Average** | **0.898713887** | **0.899669795** | **0.928996817** | **0.914056251** | **0.957081128** |

#### 4.2 Deep learning pipeline

##### 4.2.1 Input preparation

**K-mers spectral representation**: Given a genome sequence and a fixed value of $k$, we calculated a vector of size $4^k$ by counting all the occurrences of small genome fragments of length k in that genome. These small fragments are called k-mers.The frequencies of different k-mers work as a position independent feature of the genome.
**One hot vectorization**: We took the DNA sequence and converted it into overlapping k-mer sequences. Each k-mer was represented as a one-hot vector which were concatenated to produce a $(4^k * maximum\_no\_of\_k - mers)$ matrix. This representation helps to capture the information of the position of the k-mer in the DNA sequence as well. This was fed into AlexNet and InceptionNet architectures and used to predict the virulence of the sequence. We have trained InceptionNet for 150 epochs for both k=3 and k=5 length k-mers and AlexNet for 135, 100 and 100 epochs in case of k-mer length 3,4 and 5 respectively.

###### 4.2.2 Model description

**Vanilla CNN**: In this model, there are three convolutional one dimension layers with filters of size 10,20 and 30 respectively.Each of these layers are followed by a max pooling layer and a batch normalization layer. To have the final output, we have passed the flattened output through two dense layers with 500 units and 1 unit in the end.
**AlexNet**: AlexNet is a 8 layer convolutional neural network architecture which consists of Convolutional layers, Activation layers, Max Pooling layers and Dense layers. It has 5 convolutional layers, 3 max pooling layers, and 3 dense layers with one output dense-softmax layer. Conv_1, Conv_2, Conv_3, Conv_4 and Conv_5 use filters of size 11, 5, 3, 3 and 3 respectively. It is a widely used CNN architecture.
**InceptionNet**: InceptionNet consists of one or more inception modules which are blocks of parallel convolutional layers with different sized filters and max pooling layers. The result of these layers are concatenated in the end. In our model, we have used a single inception module with a fixed number of filters for each of the parallel convolutional layers. As this inception module is used twice in our model, it has computational performance-based modification. It can control the amount of reduction in the number of filters prior to convolutional layers with 3 and 5 filters respectively and the number of increased filters after max pooling.
**Experimentation Details**:

1. Training AlexNet and InceptionNet is much slower than vanilla CNN and they fill up RAM memory very quickly. So,the total number of epochs was limited between 100 and 150 at the time of training these models.

2. Since each sequence is around 29000 base pairs long, the maximum number of k-mers is also very large, around 31000. So it was filling up the RAM memory if we tried to feed this representation of all training examples at once. Hence we wrote a generator function and used it to yield a batch-size number of 2D matrices at once. Moreover each epoch takes a long time, so we have trained AlexNet for 135, 100 and 100 epochs in case of k-mer length 3,4 and 5 respectively.

### 5 Results

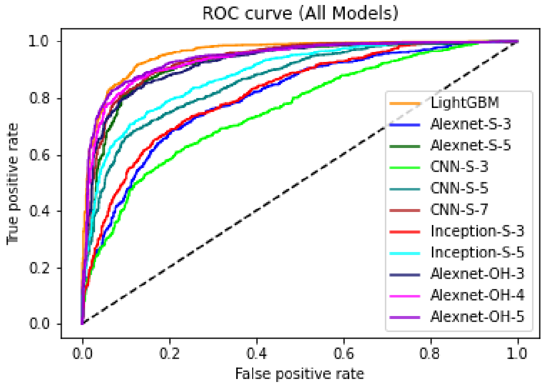


(a) ROC curve of all models

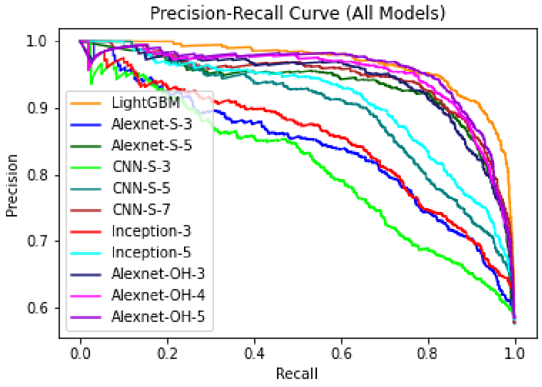


(b) Precision-Recall curve of all models

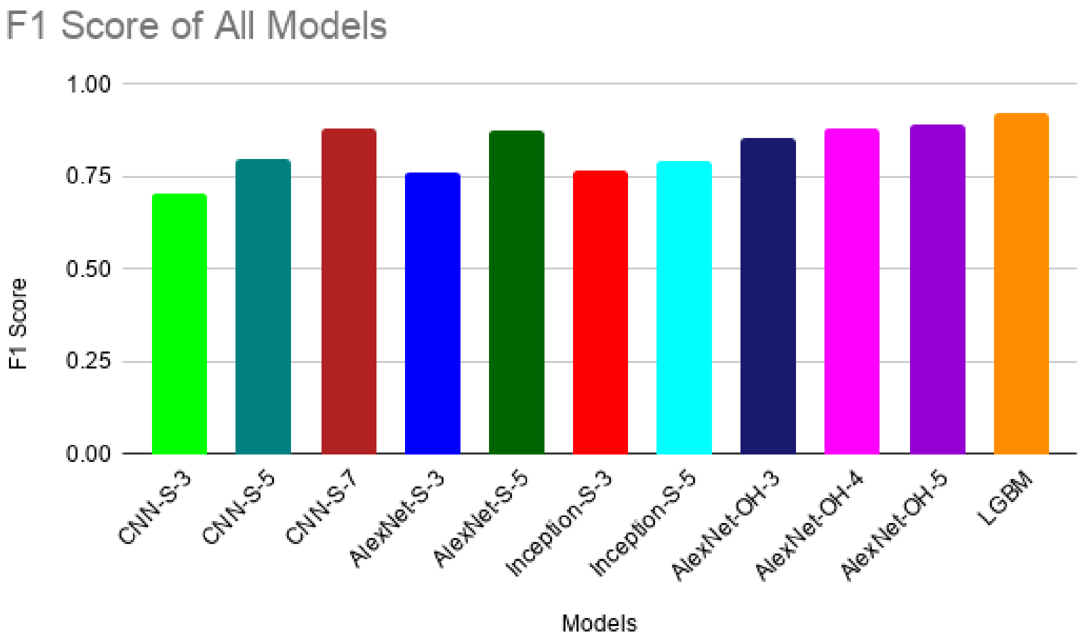


(c) F1 Score of all models

Figure 2: : ROC curve, Precision Recall curve and F1-score of all models.

Table 5: The input form, models used and accuracy, precision, recall, F1_score AUROC, AUPR are mentioned in the table. K-mer length and Epochs were additionally mentioned for the deep learning models.

| **Input Form** | **Model** | **K-mer Length** | **Epochs** | **Test Accuracy** | **Precision** | **Recall** | **F1 Score** | **AUROC** | **AUPR** |
|---|---|---|---|---|---|---|---|---|---|
| **Feature Matrix** | LightGBM Classifier | - | - | **0.8961** | **0.89585** | **0.9267** | **0.9110** | **0.95828** | **0.9670** |
| **One-Hot** | AlexNet | 3 | 135 | 0.8436 | 0.9104 | 0.8073 | 0.8557 | 0.92487 | 0.9386 |
| | | 4 | 100 | 0.8679 | 0.9085 | 0.8563 | 0.8816 | 0.93587 | 0.9480 |
| | | **5** | **100** | **0.8738** | **0.8904** | **0.8896** | **0.8900** | **0.94329** | **0.9545** |
| **Spectral K-mer Representation** | Vanilla CNN | 3 | 1000 | 0.6839 | 0.7595 | 0.6571 | 0.7046 | 0.7559 | 0.8096 |
| | | 5 | 1000 | 0.7670 | 0.7975 | 0.7961 | 0.7968 | 0.8609 | 0.8938 |
| | | **7** | **1000** | **0.8603** | **0.8563** | **0.9090** | **0.8819** | **0.9326** | **0.9429** |
| | AlexNet | 3 | 1000 | 0.7351 | 0.7907 | 0.7320 | 0.7603 | 0.8038 | 0.8402 |
| | | **5** | **1000** | **0.8550** | **0.8495** | **0.9082** | **0.8779** | **0.9213** | **0.9328** |
| | InceptionNet | 3 | 1000 | 0.7332 | 0.7626 | 0.7767 | 0.7696 | 0.8177 | 0.8517 |
| | | **5** | **1000** | **0.7844** | **0.8769** | **0.7262** | **0.7945** | **0.8828** | **0.9087** |

Table 6: Classification Results of the new severe strains.The viral strains from India and some states of the USA, before and after the cut-off date, were analyzed.

| **Countries** | **Previous Death Count/Label (24-04-2020)** | **Current Death Count/Label (25-05-2020)** | **Classifier Prediction Accuracy (Strains Submitted After the Cut Off Date)** |
|---|---|---|---|
| India | 718/Mild | 4021/Severe | >76% (3 classifiers) |
| **USA States** | | | |
| Pennsylvania | 1786/Mild | 5159/Severe | 100% (Best of 3 classifiers) |
| Maryland | 798/Mild | 2302/Severe | 100% (Best of 3 classifiers) |
| Indiana | 741/Mild | 1984/Severe | 100% (Best of 3 classifiers) |
| Illinois | 1804/Mild | 4912/Severe | >80% (3 classifiers) |
| Florida | 1045/Mild | 2251/Severe | >81.8% (3 classifiers) |

Table 7: From the classification pipeline, biologically significant features pos_19610_19614 and pos_8445_8449 are chosen using SHAP and passed down to mutation pipeline to predict mutation on. Similarly, pos_24065_24069 and pos_23825_23829 are chosen from selectKBest. The test accuracy of the mutation pipeline for these SoIs is shown in this table.

| Input Form | Method | SoI | Position | CNN-LSTM | CNN-Bidirectional LSTM |
|---|---|---|---|---|---|
| One-Hot | SHAP | 19610_19614 | 19610 | 0.7006 | 0.7006 |
| | | | 19611 | **0.7989** | **0.7989** |
| | | | 19612 | **0.7746** | 0.7745 |
| | | | 19613 | **0.7752** | **0.7752** |
| | | | 19614 | 0.7252 | **0.7255** |
| | | 8445_8449 | 8445 | 0.5496 | 0.5496 |
| | | | 8446 | 0.5496 | 0.5496 |
| | | | 8447 | 0.8733 | **0.8735** |
| | | | 8448 | 0.8974 | **0.8995** |
| | | | 8449 | **0.9269** | **0.9269** |
| | selectKBest | 24065_24069 | 24065 | 0.6264 | 0.6264 |
| | | | 24066 | 0.55337 | 0.55335 |
| | | | 24067 | **0.92325** | **0.92325** |
| | | | 24068 | **0.8743** | **0.8743** |
| | | | 24069 | **0.92475** | **0.92475** |
| | | 23825_23829 | 23825 | 0.5999 | 0.5999 |
| | | | 23826 | 0.57686 | 0.57686 |
| | | | 23827 | **0.84823** | **0.84865** |
| | | | 23828 | **0.9498** | **0.9500** |
| | | | 23829 | **0.89748** | **0.89748** |

Table 8: The mutation prediction pipeline was trained separately for Bangladesh, India and, Pakistan using Dataset A by considering the length of the concatenated time series samples up to the date of the first reported incidence of SARS-CoV-2 infected patients of that country. Later, new genome sequences were collected up to 15 May, 2020 and five new samples were generated to test the trained models. The test accuracy for Dataset A and accuracy for five new samples for various SoIs from 24065 to 24069.

| **Input Form** | **Country** | **Position** | **CNN-Bidirectional LSTM (Test accuracy / Accuracy for newly generated 5 samples)** |
|---|---|---|---|
| **One-Hot** | Bangladesh | **24065** | 0.9750/1.00 |
| | | **24066** | 0.9236/1.00 |
| | | **24067** | 0.9236/1.00 |
| | | **24068** | 1.00/1.00 |
| | | **24069** | 1.00/1.00 |
| | India | **24069** | 1.00/1.00 |
| | Pakistan | **24066** | 0.9740/1.00 |
| | | **24068** | 1.00/1.00 |
| | | **24069** | 1.00/1.00 |

Table 9: New genome sequences were again collected up to 14 December, 2020 and this time fifteen new time series samples were generated. The test accuracy for the new samples generated from new genome sequences for SoIs from 24065 to 24069 are shown in this table.

| **Input Form** | **Country** | **Position** | **CNN-Bidirectional LSTM (Accuracy for Newly Generated 15 Samples)** |
|---|---|---|---|
| **One-Hot** | Bangladesh | **24065** | 0.80 |
| | | **24066** | 0.80 |
| | | **24067** | 0.80 |
| | | **24068** | 0.867 |
| | | **24069** | 0.80 |

Table 10: We have trained our CNN-Bidirectional LSTM model for pos_8442_8446, pos_24062_24066 and pos_23822_23826 to get an improved result for the first two positions of the SoIs mentioned in Table 7. As those two positions are now at fourth and fifth position respectively, the model has predicted way better than before. So, given more information the models indeed perform better.

| **Input Form** | **Position** | **CNN-Bidirectional LSTM (Test Accuracy)** |
|---|---|---|
| **One-Hot** | **8445** | 1.0 |
| | **8446** | 1.0 |
| | **24065** | 1.0 |
| | **24066** | 1.0 |
| | **23825** | 1.0 |
| | **23826** | 1.0 |
